# The microbiota affects energy production, nitrogen excretion and sterol metabolism in mosquito larvae

**DOI:** 10.64898/2025.12.17.695059

**Authors:** Ottavia Romoli, Yanouk Epelboin, Viola Pavoncello, Frédéric Barras, Volker Behrends, Mathilde Gendrin

**Affiliations:** Microbiota of Insect Vectors Group, Institut Pasteur de la Guyane, Cayenne, French Guiana; Viruses and RNA interference Unit, Institut Pasteur, Université Paris Cité, CNRS UMR3569, Paris, France; lnserm, UA17, SPAmaz, Cayenne, French Guiana; SAMe Unit, Department of Microbiology, Institut Pasteur, Université Paris Cité, CNRS UMR6047, Paris, France; School of Medicine and Biosciences, University of West London, London, W5 5RF; Faculty of Medicine, Imperial College London, London W12 0HS

**Keywords:** Mosquito, *Aedes aegypti*, larval development, metabolome, GC-MS, microbiota, transient colonization, nitrogen metabolism, TCA cycle, cholesterol, fatty acids, beta-oxidation

## Abstract

Mosquito larvae rely on a living microbiota for normal development because the microbiota supplies essential nutrients, particularly vitamins. Beyond vitamin provision, transcriptomic data suggest that the microbiota also supports other key nutritional processes. Here, we explored these roles by conducting a metabolomics analysis on *Aedes aegypti* third instar larvae following microbiota depletion. We sampled larvae and dissected guts 12- and 20-hours post-decolonization and analysed methanol-soluble metabolites using untargeted gas chromatography–mass spectrometry. Our findings reveal a pronounced impact of gut microbial presence on several metabolites involved in the tricarboxylic acid cycle and the uricolytic pathway. Germ-free larvae also had a lower quantity of cholesterol in guts and their long-chain fatty acid profile was altered in guts and whole larvae. Sterols, including cholesterol, are essential precursors for the production of the moulting hormone 20-hydroxyecdysone. We therefore tested how supplementing exogenous cholesterol affects the development of germ-free larvae. The effects proved to be highly concentration-dependent, ranging from a marginally significant increase in successful development to adulthood at low concentrations to a pronounced developmental impairment at higher concentrations. Moreover, bacteria deficient in fatty acids beta-oxidation had a significantly lower ability to support larval development. Together, the observed alterations suggest that microbiota-deprived larvae exhibit a downregulation of metabolic processes related to energy production, nitrogen excretion and sterol metabolism, likely due to the absence of microbiota-derived vitamins essential for these central metabolic functions.

**Importance:** Mosquito larvae depend on gut microbiota for normal growth because microbes supply essential nutrients, particularly B vitamins. To explore microbial roles beyond vitamin provision, we analysed metabolic changes in *Aedes aegypti* larvae after microbiota removal using gas chromatography-mass spectrometry. Germ-free larvae exhibited decreased metabolites associated with the tricarboxylic acid cycle and uricolytic pathway, indicating a general slowdown in metabolic activity and nitrogen waste processing. Additionally, the absence of a microbiota affected cholesterol and fatty acid metabolism. To validate these findings, we found that supplementing germ-free larvae with low levels of cholesterol modestly improved their development. In contrast, larvae colonized with bacteria deficient in fatty acid metabolism exhibited significantly reduced developmental success. Overall, the findings show that removing the microbiota downregulates key metabolic pathways related to energy production, nitrogen excretion, and sterol metabolism, highlighting that bacterial vitamins and fatty acid degradation are vital for mosquito larval development and successful transformation into adults.

## Introduction

Mosquitoes are holometabolous insects undergoing an aquatic larval development and a terrestrial adult phase. Larval development is the life stage in which the mosquito acquires the totality of energy and elements to form a mature adult. These two distinct developmental stages are characterized by intrinsically different environments and nutritional requirements. During adulthood, mosquitoes fly, feed on plant nectar, and females generally ingest blood, as they require this nutritionally rich diet for egg production. During larval development, mosquitoes feed on organic debris and uptake microorganisms that colonize their gut and critically support their development^1,2^. Some microorganisms are transmitted from larvae to adults, while others are taken up by adults from their environment. The composition of this microbiota at both larval and adult stage significantly influences physiology, consequently affecting vector competence to pathogens, fecundity and lifespan^3–9^. Importantly, microbiota composition during larval development has been reported to influence adult physiology without influencing adult microbiota composition, indicating that host-microbe interactions that happen during larval development matter on pathogen transmission^8,10^. Hence, the mosquito microbiota has a strong influence on its host throughout the aquatic and aerial phases of its life cycle.

Focusing on larvae, microbes appear to be required for normal development of the host. In the absence of microorganisms, it is only possible to rear larvae in the dark with a special liver and yeast extract-based^11^ diet or with a synthetic medium that includes vitamins, salts and amino acids^12^. In contrast, germ-free larvae kept on conventional day/night cycles, whatever the diet, are blocked in their first instar until microbes are added to the environment^2,11,12^. Such microbes are not particularly specific, several families of bacteria as well as yeasts and algae have been found to support larval development, and differences between microbial strains in development success depend on several factors such as virulence and colonization success^13–15^. The non-specificity in the taxonomic entities favouring development suggested that microbial support is linked to relatively common constituents, and recent work indeed showed that bacterial support on development notably relies on the provision of B vitamins to their host. On one hand, our transcriptomic analysis on germ-free vs colonized third instar larvae detected an upregulation of genes of the folate (B9 vitamin) biosynthesis pathway, which is incomplete in mosquitoes, and of two folate transporters^4^. We observed that, after turning germ-free during the third instar, most larvae did not develop to adulthood, but development success increased from 12 % to 52 % upon folate supplementation. On the other hand, Wang *et al* defined a diet allowing to support development of axenic larvae and showed that an essential dietary compound was light sensitive; they identified riboflavin (B2 vitamin) as an essential dietary requirement for axenic larvae^12^. They also found that pyridoxin (B6 vitamin), and to a lesser extent thiamine (B1 vitamin) and folate, were required for larval development, while pantothenate (B5 vitamin), nicotinic acid (B3 vitamin) and biotin (B7 vitamin) were not. Our recent work indicates a positive impact of biotin when supplemented with folate, but that biotin also has a toxic effect at higher concentrations^16^.

Considering the importance of bacterial colonization on mosquito metabolism, we decided to study the metabolome of *Aedes aegypti* larvae colonized or not with bacteria, using a similar experimental strategy as for our previous study. It consists in colonizing germ-free first instar larvae with a bacterial strain auxotroph for meso-diaminopimelic acid and D-alanine, two bacteria-specific amino acids^17^. Bacteria are lost by most larvae by 12 h after transfer into an environment devoid of both amino acid supplements. Here, we report our data after gas-chromatography-mass spectrometry (GC-MS) analysis on gut and whole larvae samples collected during the second half of the third instar, i.e. 12 h and 20 h after larval transfer to the bacteria- and supplement-free environment. We found that larvae deprived of their microbiota show an increase of metabolites involved in the citric acid (TCA) cycle, a decrease of cholesterol and metabolites involved in the uricolytic pathway and a general perturbation of unsaturated fatty acid metabolism. When supplementing cholesterol to germ-free larvae, we detect a marginally significant increase of development to adulthood at low concentrations, suggesting an involvement of this metabolite in mosquito larval development. Additionally, we demonstrate that bacteria contribute to larval development by degrading and providing fatty acids to the host.

## Results

### Absence of microbiota in mosquito larvae alters central metabolic pathways, including the TCA cycle and uricolytic pathway

Our previous study on the role of the microbiota in *Ae. aegypti* larvae identified specific transcriptional signatures indicating a strong involvement of bacteria on mosquito metabolism during larval development^4^. Thus, we decided to assess the impact of bacterial decolonization on the larval metabolome, using an experimental set up similar to our previous work, i.e. focusing on axenic third instar larvae produced via transient colonization with *Escherichia coli* HA416 auxotrophic bacteria. Our control consisted of larvae colonized with the parental WT strain *E. coli* HS. We sampled whole larvae or dissected guts 12 h and 20 h after transfer of early third instar larvae into water that contained fresh sterile food but no bacteria (Figure 1). As per our previous transcriptomic study, dissected guts were sampled to investigate processes linked to nutrition, while whole larvae were used to study whole body metabolism. After methanol extraction, we performed a GC-MS analysis of the gut and larval metabolome. This analysis allowed us to specifically quantify 80 metabolites notably including 17 amino acids, 11 fatty acids, 7 sugars, and other metabolites implicated in diverse pathways (Table S1).

**Figure 1.**
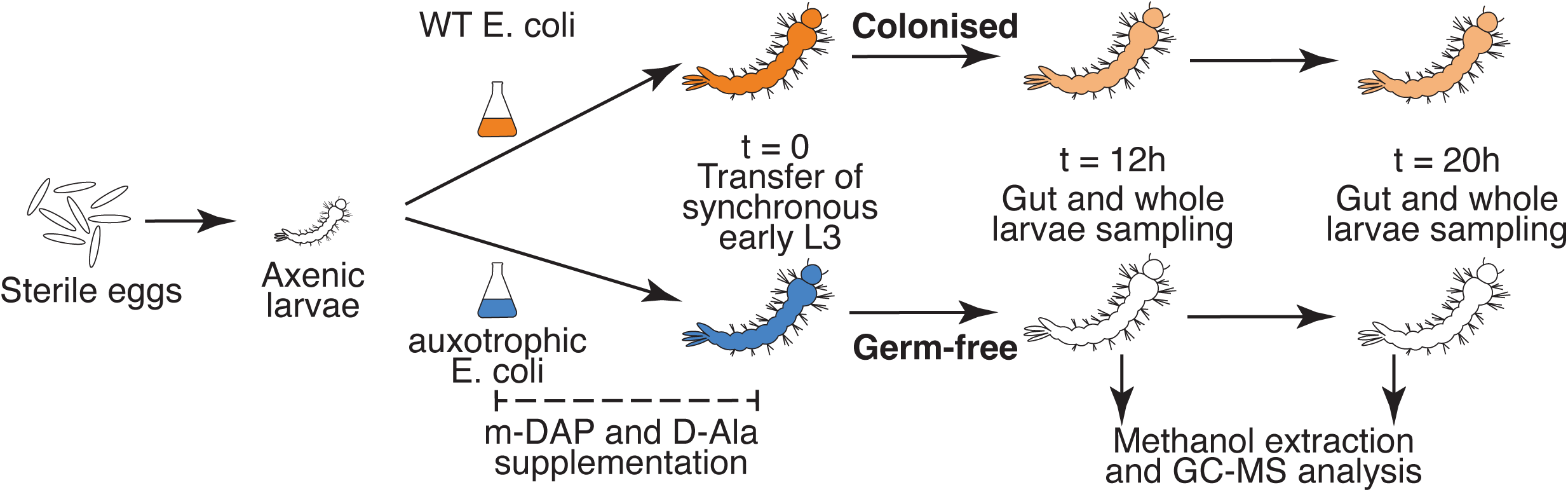
Experimental set-up. Axenic *Aedes aegypti* larvae were obtained by egg surface sterilization. To support larval development, larvae were either colonized with wild-type (WT) *Escherichia coli* (orange) or supplemented with an isogenic *E. coli* strain auxotroph (AUX) for the bacteria-specific amino acids meso-diaminopimelic acid (m-DAP) and D-alanine (D-Ala, blue), which were added to the larval water. At the onset of the third instar, both WT- and AUX-colonized larvae were transferred to sterile water containing only sterile fish food (i.e., no bacteria or amino acids). This allowed to achieve bacterial decolonisation from AUX-carrying larvae and obtain germ-free larvae, while WT-carrying larvae were still colonised by bacteria. Larvae were collected at 12- and 20-h post-transfer for metabolite extraction, from pools of dissected guts and whole larvae. Metabolomic profiles were analysed by gas chromatography–mass spectrometry (GC-MS). The experiment was repeated in four independent replicates, collecting 50-60 individuals per replicate and sample type.

Principal component analysis of all metabolites across all samples revealed a clear separation between gut and larva samples along the first principal component (Figure 2). Tissue type emerged as the primary explicatory variable driving the differences in metabolome composition, while bacterial colonization had only a marginal impact and the interaction between colonization and time failed to provide a solid explanation for the variation of metabolite levels (PERMANOVA on tissue, based on Euclidean distance, F = 22.7, *p* = 0.001; PERMANOVA on colonization: F = 2.0, *p* = 0.085; PERMANOVA on colonization*tissue: F= 10.1, *p* = 0.001; PERMANOVA on colonization*time: F = 1.2, *p* = 0.25, Table S2).

**Figure 2.**
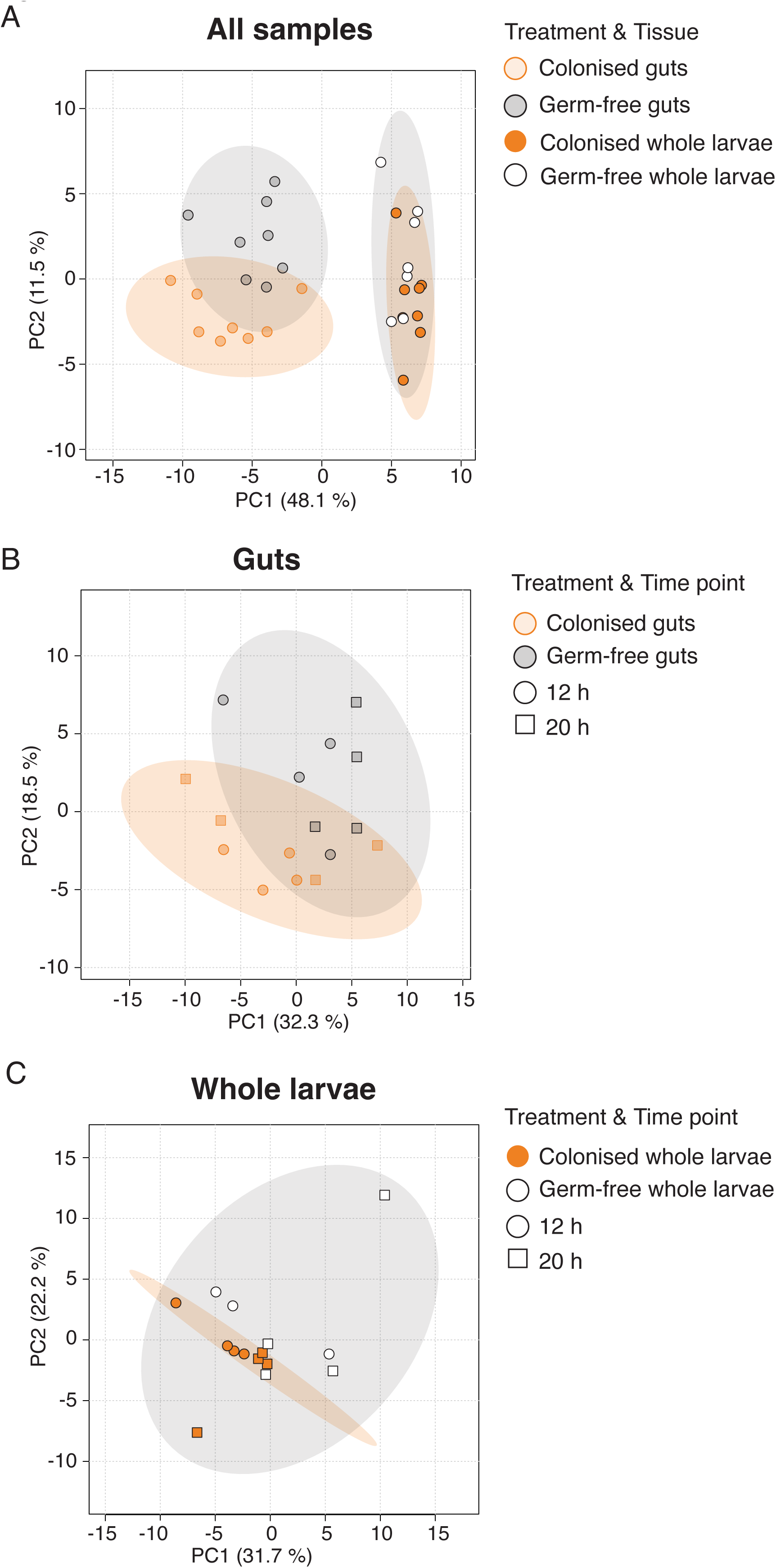
Principal Component Analysis (PCA) of gut and larval metabolomes. (**A**) PCA plot showing a clear separation along PC1 between gut (transparent symbols) and whole larva (solid symbols) metabolomes (PERMANOVA tissue: F = 22.7, *p* = 0.001). Samples are also partially grouped by colonization status (orange: colonized, white/grey: germ-free; PERMANOVA colonization: F = 2.0, *p* = 0.085). (**B**) PCA of gut metabolomes, with partial separation by colonization but not by time-point (circles: 12 h; squares: 20 h; PERMANOVA colonization: F = 2.6, *p* = 0.01; time-point: F = 1.1, *p* = 0.39). (**C**) PCA of whole larva metabolomes, showing partial separation between colonized and germ-free samples and between 12 h and 20 h samples (PERMANOVA colonization: F = 1.9, *p* = 0.039; time: F = 2.1, *p* = 0.023). In each panel, grey and orange ellipses display 95 % confidence regions of germ-free and colonised samples, respectively.

Differences linked to bacterial colonization were significant when focusing on gut and whole larva samples separately (Figure 2B – PERMANOVA: F = 2.6, *p* = 0.014 on guts; Figure 2C – PERMANOVA: F = 1.8, *p* = 0.039 on whole larvae). Interestingly, time points significantly discriminated whole larva samples (PERMANOVA: F = 2.1, *p* = 0.023), while they did not affect the metabolome structure of gut samples (PERMANOVA: F = 1.0, *p* = 0.39). This contrasts with transcriptomic data, where sampling time points had a strong impact both in guts and larvae^4^. Therefore, in the following analyses, we prioritized identifying differences between germ-free and colonized samples over those associated with time points.

Gut and larval samples exhibited distinct metabolite profiles, suggesting the compartmentalization of specific metabolic pathways inside or outside the gut. Specifically, gut metabolome was enriched in fatty acids (such as lauric acid/C12:0 or palmitic acid/C16:0), while larval samples showed high levels of amino acids (including methionine, tyrosine, and lysine) and monoacylglycerols (such as C16:0-glycerol, Figure 3).

**Figure 3.**
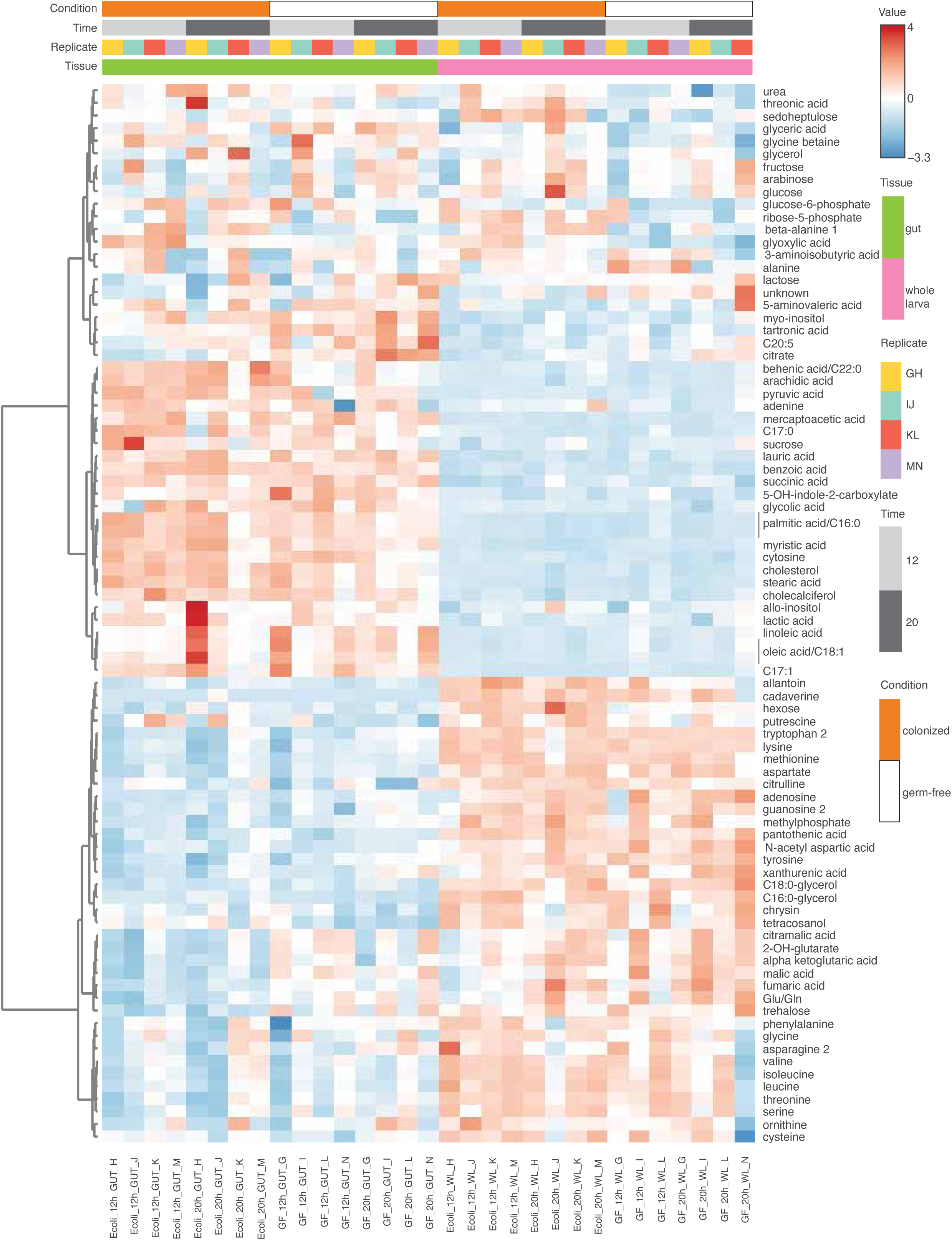
Heatmap of gut and larval metabolomes. Heatmap displaying metabolite intensities across samples, organized by colonization status, time point and tissue type. Hierarchical clustering based on Euclidean distances was performed using Ward’s method. Two primary clusters emerge, reflecting distinct metabolite profiles between gut and whole-larva samples.

Focusing on the impact of bacterial colonization, we observed that 28 and 20 metabolites were significantly affected in guts and whole larvae, respectively, with t-test thresholds at *p* = 0.1 and |fold change| = 1.2 (Figure 4, Figure S1, Table S2). Levels of several metabolites of the tricarboxylic acid (TCA) cycle were increased in germ-free conditions – citrate and malate in guts and larvae and alpha-ketoglutarate and fumarate in guts (Figure 4). Germ-free gut samples also showed a reduced quantity in some lipids and fatty acids (behenic acid (C22:0), arachidic acid (C20:0), palmitic acid (C16:0), monopalmitoylglycerol (C16:0-glycerol)), or sterols (tetracosanol, cholesterol). On the contrary, other fatty acids were more abundant in germ-free whole larvae, namely linolenic acid (C18:3), linoleic acid (C18:2), oleic acid (C18:1), and myristic acid (C14:0), while metabolites of the nitrogen homeostasis pathway (urea, glyoxylic acid and allantoin) and sugars (hexose, ribose-5-phosphate) were depleted (Figure 4, Figure S1, and Table S2). Germ-free guts and larvae were also deprived in beta-alanine, a metabolite involved in the biosynthesis of pantothenate (B5 vitamin).

**Figure 4.**
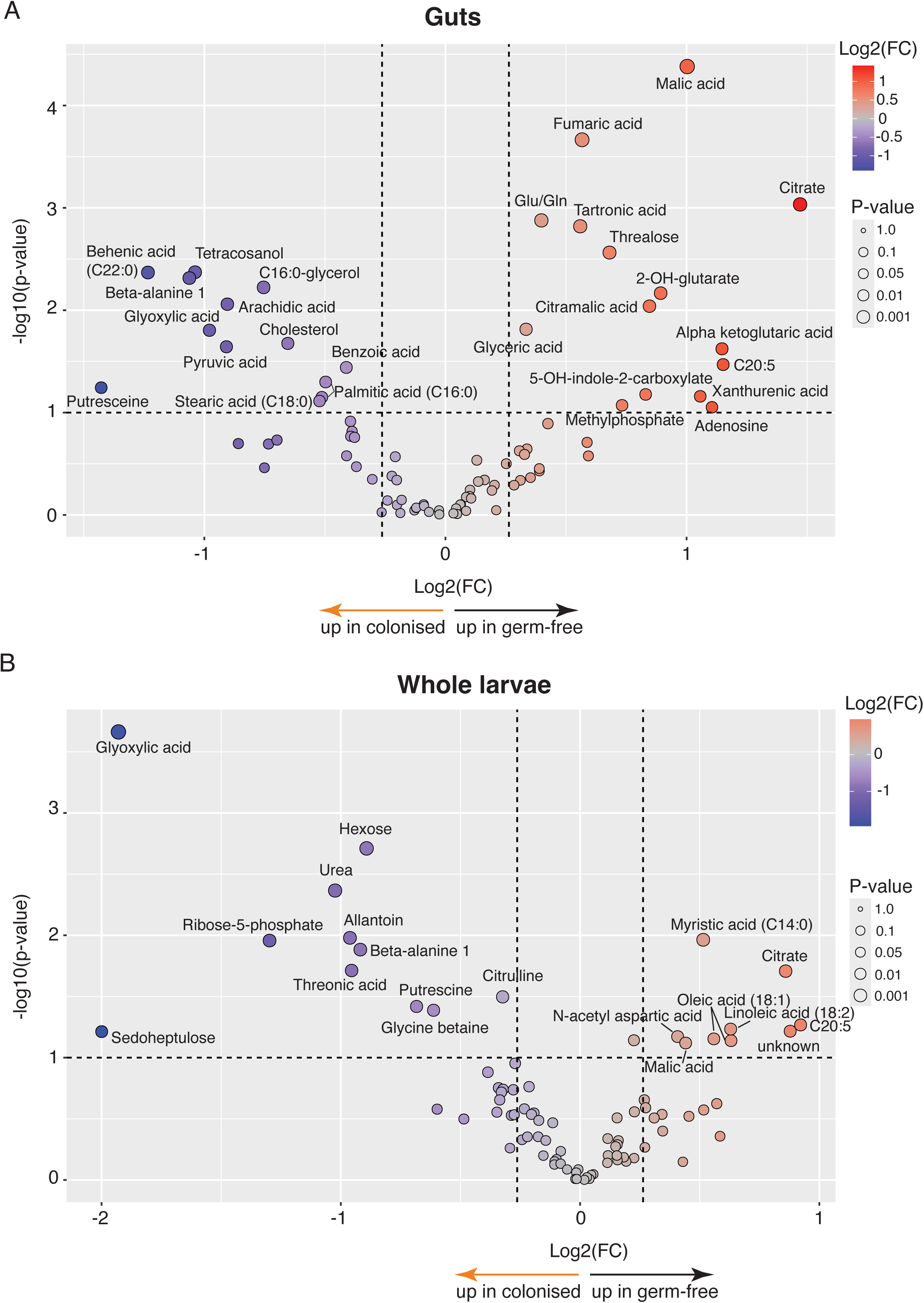
Metabolites significantly affected by bacterial colonization in gut and larval samples. Volcano plots showing differential metabolite abundances between colonized and germ-free conditions in gut (**A**) and whole-larva (**B**) samples. Metabolites enriched in colonized samples are indicated by orange arrows; those enriched in germ-free samples by black arrows. A fold change threshold of 1.2 and a t-test *p* value threshold of 0.1 were applied. Both fold changes and *p* values are log-transformed for visualization.

Finally, differentially abundant metabolites were subjected to a pathway enrichment analysis, which indicated a strong effect of bacterial colonization on (i) TCA cycle (guts: *p* < 0.001, 5 hits, i.e. 5 affected metabolites; larvae: *p* = 0.022, 2 hits), (ii) glyoxylate and dicarboxylate metabolism (guts: *p* < 0.001, 7 hits; larvae: *p* = 0.002, 3 hits), (iii) alanine, aspartate and glutamate metabolism (gut: *p* < 0.001, 6 hits; larvae: *p* = 0.041, 2 hits), (iv) arginine biosynthesis (gut: *p* < 0.001, 4 hits; larvae: *p* = 0.011, 2 hits) and (v) synthesis of unsaturated fatty acids (guts and larvae: *p* < 0.001, 4 hits, Table 1). Additionally, other pathways connected to the TCA cycle were impacted by bacterial colonization in the gut, such as pyruvate metabolism (*p* = 0.003, 3 hits) and arginine and proline metabolism (*p* < 0.001, 4 hits).

**Table 1.** Pathway enrichment analysis of metabolites affected by bacterial colonization in gut and larva samples. For each pathway, the following elements are indicated: total number of metabolites involved in the pathway, expected number of metabolites found in the pathway, number of metabolites affected by bacterial colonization, raw p value and false discovery rate (FDR). Pathways with an FDR < 0.05 are highlighted in grey.

| Pathway | Total metabolites | Expected metabolites | Hits | Raw *p* | FDR |
| --- | --- | --- | --- | --- | --- |
| **Gut (colonized vs germ-free)** |  |  |  |  |  |
| Glyoxylate and dicarboxylate metabolism | 31 | 0.40 | 7 | 4.3E-08 | 3.4E-06 |
| Alanine, aspartate and glutamate metabolism | 28 | 0.36 | 6 | 6.7E-07 | 2.7E-05 |
| Citrate cycle (TCA cycle) | 20 | 0.26 | 5 | 3.0E-06 | 7.9E-05 |
| Arginine biosynthesis | 14 | 0.18 | 4 | 1.9E-05 | 0.00038 |
| Arginine and proline metabolism | 36 | 0.47 | 4 | 0.00094 | 0.013 |
| Biosynthesis of unsaturated fatty acids | 36 | 0.47 | 4 | 0.00094 | 0.013 |
| Nitrogen metabolism | 6 | 0.08 | 2 | 0.0023 | 0.027 |
| Pyruvate metabolism | 23 | 0.30 | 3 | 0.0028 | 0.028 |
| Glycine, serine and threonine metabolism | 33 | 0.43 | 3 | 0.0080 | 0.071 |
| Butanoate metabolism | 15 | 0.20 | 2 | 0.015 | 0.11 |
| Tyrosine metabolism | 42 | 0.55 | 3 | 0.016 | 0.11 |
| Glycerolipid metabolism | 16 | 0.21 | 2 | 0.017 | 0.12 |
| Glutathione metabolism | 28 | 0.36 | 2 | 0.050 | 0.28 |
| Lipoic acid metabolism | 28 | 0.36 | 2 | 0.050 | 0.28 |
| **Larvae (colonized vs germ-free)** |  |  |  |  |  |
| Biosynthesis of unsaturated fatty acids | 36 | 0.42 | 4 | 0.00061 | 0.049 |
| Glyoxylate and dicarboxylate metabolism | 31 | 0.36 | 3 | 0.0049 | 0.20 |
| Arginine biosynthesis | 14 | 0.16 | 2 | 0.011 | 0.29 |
| Citrate cycle (TCA cycle) | 20 | 0.23 | 2 | 0.022 | 0.43 |
| Pentose phosphate pathway | 23 | 0.27 | 2 | 0.028 | 0.45 |
| Alanine, aspartate and glutamate metabolism | 28 | 0.33 | 2 | 0.041 | 0.51 |
| Purine metabolism | 70 | 0.82 | 3 | 0.045 | 0.51 |

Overall, these findings highlight a significant influence of the microbiota on the urea synthesis pathway in whole larvae (Figure 5) and on the TCA cycle in guts (Figure 6). Specifically, several TCA cycle metabolites were more abundant in germ-free guts. Consistent with this, transcriptomic data show downregulation of multiple genes encoding key TCA cycle enzymes in germ-free guts (Figures 6,^4^).

**Figure 5.**
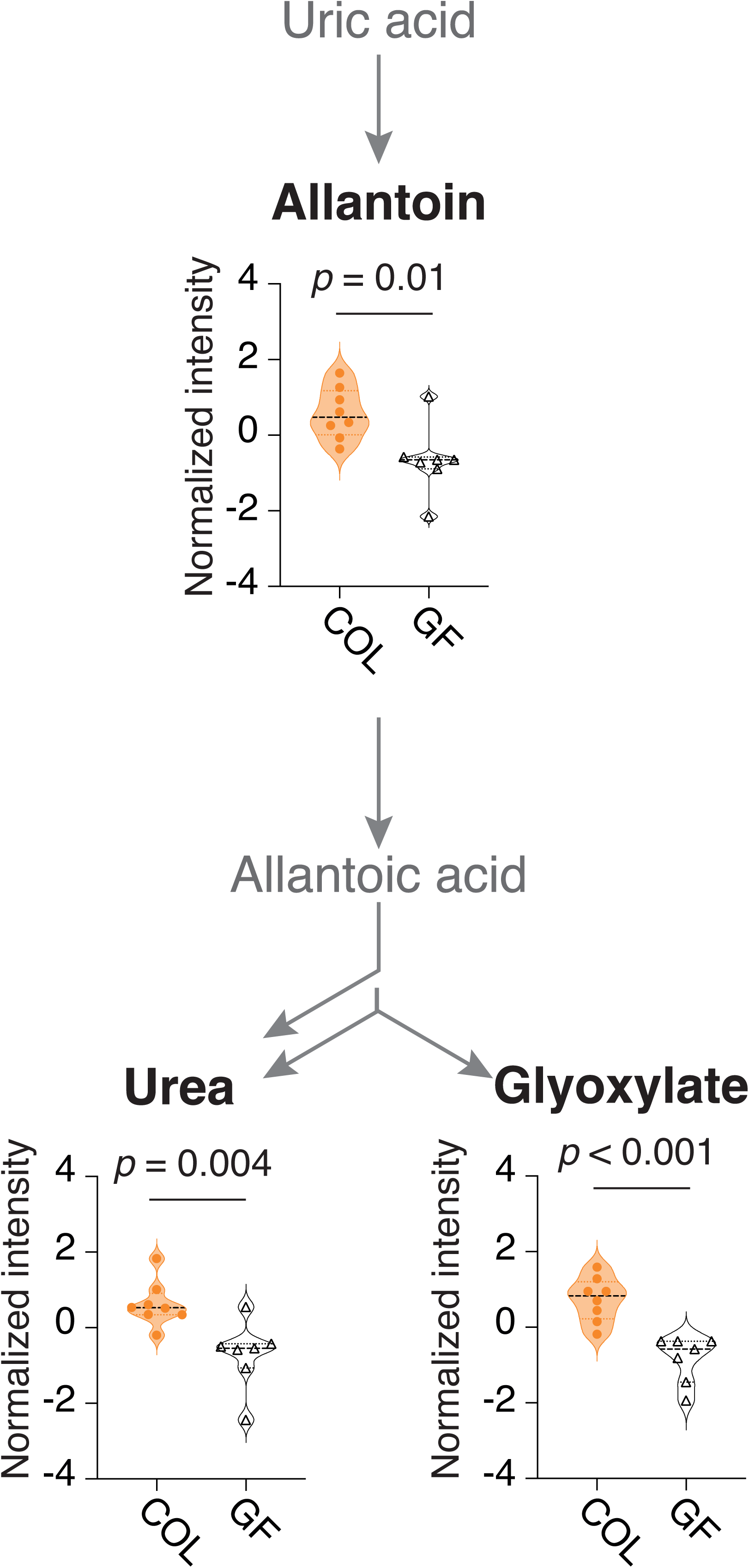
Uricolytic pathway metabolites affected by bacteria colonization in mosquito larvae. Key reactions of the uricolytic pathway are shown: uric acid is oxidized to allantoin, which is subsequently converted to allantoic acid. Allantoic acid is further degraded into two urea molecules and one glyoxylate molecule. Metabolites significantly impacted by bacterial colonization in whole larvae (based on t-test) are highlighted in bold; metabolites not detected in our metabolomic dataset are shown in grey. For detected metabolites, violin plots display normalized intensities in colonized (COL, orange) and germ-free (GF, white) samples. Median values are indicated by black dashed lines, and quartiles by orange or black dotted lines.

**Figure 6.**
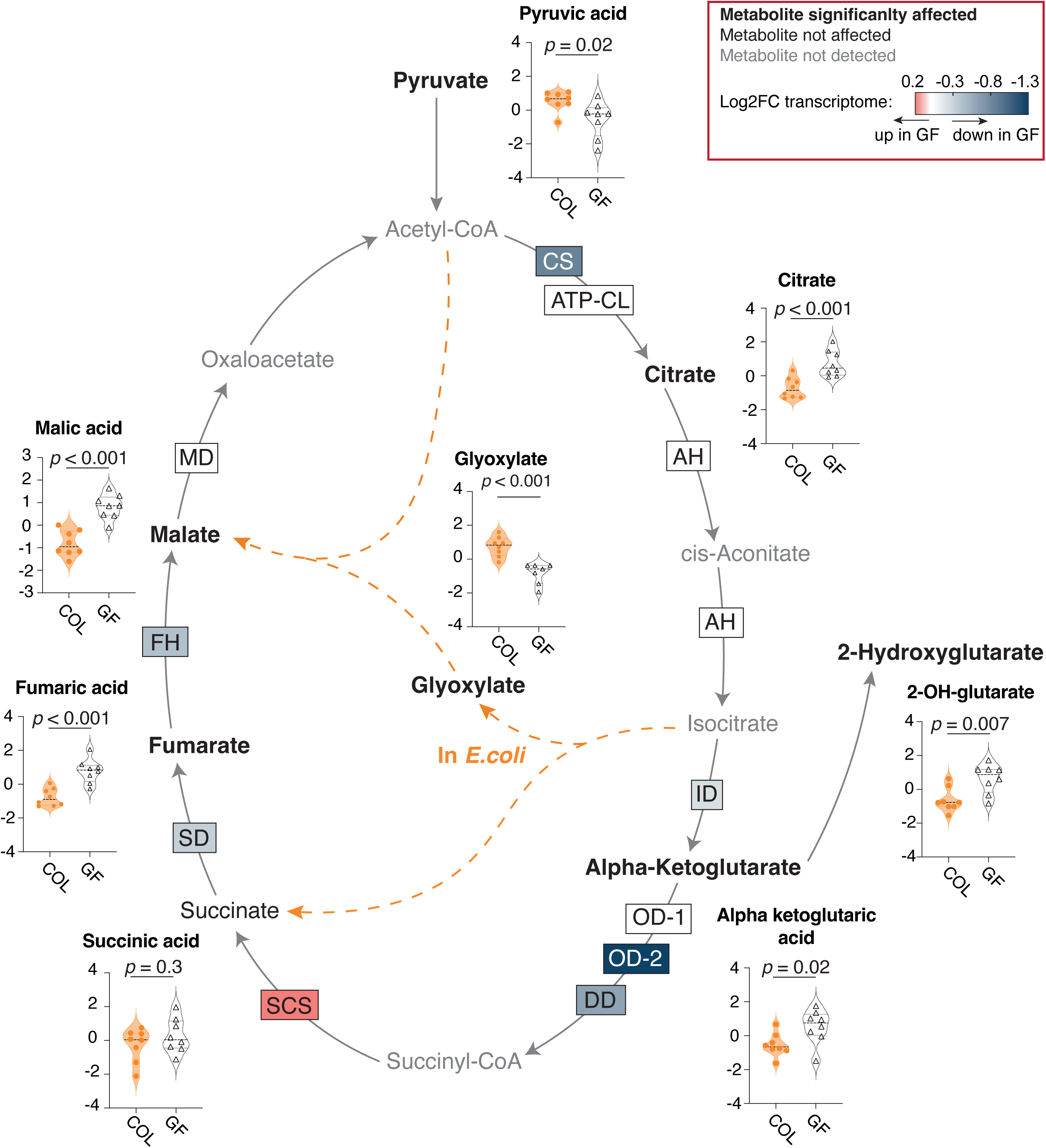
Tricarboxylic acid (TCA) cycle metabolites affected by bacteria colonization in larval guts. Key metabolites and enzymes of the TCA cycle are shown. Metabolites significantly altered by bacterial colonization in larval guts (based on t-test) are highlighted in bold, while those not detected in our metabolomic dataset are shown in grey. Enzyme boxes backgrounds are color-coded according to GF/WT log_2_ fold change values from our previous transcriptomic analysis^4^, averaged across the 12 h and 20 h time points. For detected metabolites, violin plots display normalized intensities in colonized (COL, orange) and germ-free (GF, white) samples. Median values are indicated by black dashed lines, and quartiles by orange or black dotted lines. Enzyme names: CS: Citrate Synthase; ATP-CL: ATP Citrate (pro-S)-Lyase; AH: Aconitate Hydratase; ID: Isocitrate Dehydrogenase; OD-1: 2-Oxoglutarate Dehydrogenase (E1); OD-2: 2-Oxoglutarate Dehydrogenase (E2); DD: Dihydrolipoyl Dehydrogenase; SCS: Succinyl-CoA-Synthetase; SD: Succinate Dehydrogenase; FH: Fumarate Hydratase; MD: Malate Dehydrogenase.

### Cholesterol affects development of germ-free larvae in a dose-dependent fashion

Our dataset comprises larval and bacterial metabolites whose level are controlled by multiple pathways, thereby rendering the source of their variation difficult to assess. In contrast, cholesterol, which we found to be depleted in germ-free guts (Figure 7A), is exclusively derived from the diet^18^. Besides its conventional role in membrane homeostasis, cholesterol is crucial for insect development as it is the initial metabolite of the 20-hydroxyecdysone biosynthesis pathway, the insect hormone critical to initiate moulting and metamorphosis. Two peaks of ecdysteroids are observed, one during the fourth (and last) larval instar and a stronger one at the beginning of metamorphosis^19^. When larvae become germ-free at the start of the third instar, most fail to complete their development, often stalling at the fourth instar or dying^4,16^. We hypothesized that a supplementation in cholesterol may increase the ability of larvae to produce ecdysteroids, hence, to go through the fourth instar development checkpoint. In two sets of experiments, we tested the impact of four cholesterol concentrations on late larval development and metamorphosis of larvae that were decolonized as third instar. Firstly, we added 35.6 and 106 µg/mL, two concentrations that frame those used by Singh and Brown (i.e. 39 µg/mL), to support larval development of germ-free *Ae. aegypti*^20^. Cholesterol had a negative impact on larval development at such concentrations (Figure 7B , generalised linear mixed model (GLMM) on developed individuals: *p* < 0.001, Table S2). Both concentrations significantly impacted the proportion of immatures dead or blocked in their development (GLMM on developed individuals: 35.6 µg/mL, *p* = 0.007; 106 µg/mL, *p* < 0.001, Table S2). High cholesterol diet is known to have negative impacts on animal health; hence these concentrations added to the cholesterol already present in the food used to feed larvae probably mimicked the impact of a high cholesterol diet. We therefore lowered the doses of cholesterol supplementation (0.36 µg/mL and 3.6 µg/mL) and tested their impact on larval development. Overall, cholesterol supplementation had a positive, albeit only marginally significant effect on developmental success (Figure 7C, GLMM on developed individuals: *p* = 0.074, Table S2). Considering each concentration separately, the effect was again marginally significant at 0.36 µg/mL (GLMM on the proportion of developed adults: 0.36 µg/mL, *p* = 0.09; 3.6 µg/mL, *p* = 0.11). We more specifically noticed that the proportion of mosquitoes dying during metamorphosis was lower in the presence of low cholesterol supplementation, particularly at 3.6 µg/mL (Figure 7C, GLMM on the proportion of dead pupae, *p* = 0.074; 3.6 µg/mL: *p* = 0.040; 0.36 µg/mL: *p* = 0.27, Table S2).

**Figure 7.**
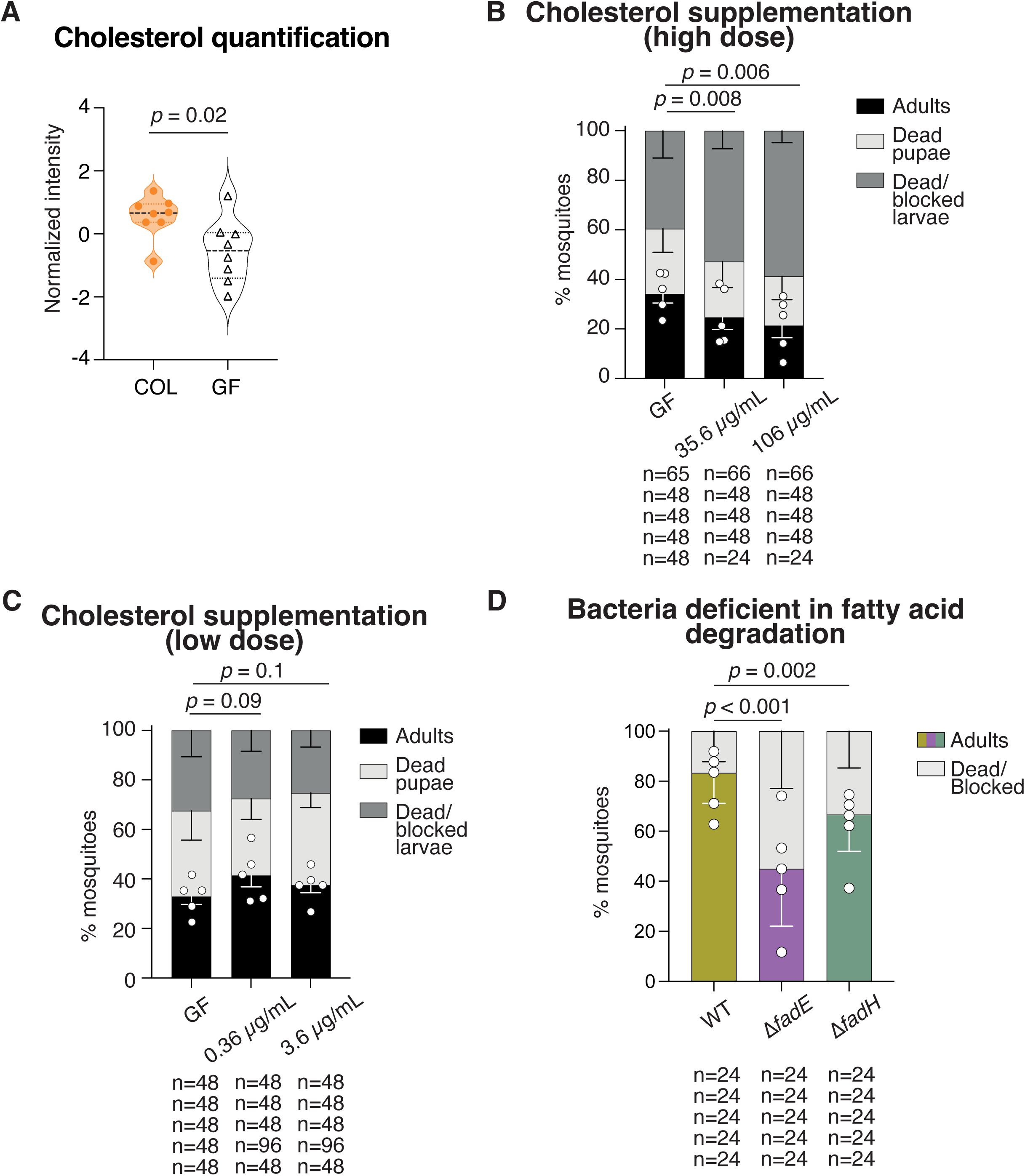
Effect of cholesterol supplementation and bacterial fatty acid degradation on mosquito larva development. (**A**) Normalized cholesterol intensities in colonized (COL, orange) and germ-free (GF, white) larval guts, compared using a t-test. Median values are indicated by black dashed lines, and quartiles by orange or black dotted lines. (**B**-**C**) Developmental outcomes under germ-free conditions with or without cholesterol supplementation. Bar plots show the proportions of adults (black) and developmentally blocked or dead larvae (grey) following supplementation with either high (**B**) or low (**C**) cholesterol doses. (**D**) Developmental outcomes of gnotobiotic larvae colonized by wild-type *E. coli* (WT, khaki) or *E. coli* mutants in *fad*E (purple) and *fad*H (green) genes. Bar plots show the proportions of adults (black, purple or green) and developmentally blocked or dead larvae (grey). All plots show results from five independent replicates. The number of individuals analysed in each replicate is indicated below the plots. Statistical significance on the proportion of adults was assessed using a generalized linear mixed model (GLMM) using the proportion of adults and the cholesterol concentration or bacterial strain as fixed factors, and the replicate as a random factor.

Overall, while our data with lower concentrations do not allow to draw strong conclusions, they suggest that minute supplementation of cholesterol modestly promote development, potentially by supporting lipid and/or ecdysteroid metabolism particularly during metamorphosis, while high concentrations mimic the negative impact associated with a high-cholesterol diet.

### Degradation of fatty acids by gut bacteria is critical for mosquito larval development

Finally, we decided to investigate the biological relevance of the variation in fatty acid levels observed in the metabolomic analysis. To test if fatty acid degradation by bacteria is important to larval development, we tested *fadE* and *fadH E. coli* mutants, which are deficient in fatty acid degradation. FadE is an acyl-CoA dehydrogenase that catalyses the first step of β-oxidation of fatty acids. FadH provides the β-oxidation entry point for degradation of unsaturated fatty acids^21,22^. An *E. coli fadE* mutant was previously shown to fail larval colonization in gnotobiotic *Ae. aegypti* when inoculated at 10^6^ CFUs/mL but could support larval growth when supplemented at higher doses^15^. In our hands, when *fadE* and *fadH* mutants were provided at low doses to germ-free larvae (i.e. ∼10^3^ CFUs/mL), bacteria could be recovered from both larval water (Figure S2A) and fourth instar larvae (Figure S2B). This indicates that these bacterial mutants did not show larval colonization defects in our experimental conditions. When provided at this low dose, we observed that both mutants had a significant impact on larval development, reducing the overall percentage of mosquitoes completing their development (Figure 7D). Specifically, *fadE*-deficient bacteria were significantly less able to support development vs *fadH*-deficient mutants, suggesting the importance of fatty-acid degradation for larval development (GLMM on developed individuals: WT vs Δ*fadE*: *p* < 0.001, WT vs Δ*fadH*: *p* = 0.002, Figure 7D). Using high bacterial concentrations, we did not detect any difference in development success, indicating that the effect of *fadE* and *fadH* on development is not linked to any kind of negative impact of these mutants on fitness, but rather reflects metabolic restriction (Figure S2C). Overall, our data indicate that fatty acid degradation by gut bacteria is important for larval development.

## Discussion

In this study, we investigated metabolic consequences of bacterial colonization in third-instar mosquito larvae. Our data notably point to microbiota-related regulation of several metabolites of the TCA cycle and urea synthesis. Additionally, the presence of microbes in the gut significantly affected the abundance of several lipids, fatty acids and cholesterol. We further show that dietary complementation in cholesterol and bacterial fatty acid degradation impact larval development.

The present study complements two previous studies of our team and will be discussed in light of these published data. Firstly, we performed a transcriptomic study on colonized vs germ-free larvae and dissected guts^4^. This study revealed stronger variations between colonized and germ-free conditions than the current metabolomic dataset, which we suspect roots in a small modification in experimental setup. We only transferred auxotrophic *E. coli*-carrying larvae into a fresh sterile larval environment for RNA-sequencing, while we now transferred both larvae carrying wild-type and auxotrophic *E. coli*. This adjustment minimized metabolic variations caused by differences in nutrient processing time (i.e., from the first instar vs only after decolonisation), which is crucial for a metabolomic study. This revised setup however transiently reduced microbiota numbers in colonized larvae during the initial hours post-transfer, which affects mosquito development even though microbial populations had rebounded in colonized larvae by the time of sampling^4^. Hence, the current study potentially missed some finely microbiota-regulated metabolites, but offers stringent control of the dietary context.

Secondly, we performed a metabolomic study of adult *Anopheles* mosquito guts, reared under conventional conditions and treated or not with antibiotics^23^ . While the specific metabolites identified in adults gut^23^ and in the present analysis differ due to a quality control step that retained only reproducible signals across each dataset, some shared metabolic patterns were comparable. Both metabolomics studies detected elevated levels of several TCA cycle intermediates in the (relative) absence of gut bacteria: citrate, fumarate, alpha-ketoglutarate, and malate in germ-free larval guts, and citrate and isocitrate in antibiotic-treated adult guts. In parallel, both datasets showed reduced levels of various sugars upstream of glycolysis in bacteria-free conditions^23^. Interestingly, several enzymes of the TCA cycle were down-regulated in transcriptome data on germ-free larvae, including the rate-limiting step of isocitrate dehydrogenase. Hence, the observed enrichment in TCA metabolites may result from a decreased flux of metabolites through this cycle. Thus, bacterial colonization appears to influence central carbon metabolism across developmental stages with an overall slowdown of the TCA cycle and thus a potential decrease of ATP production in cells.

Considering nitrogen excretion, we detected urea rather than uric acid as the prominent ammonia waste product in larvae, which aligns with previous reports of low uric acid levels in third-instar larvae compared to adults^24^. In *Ae. aegypti*, urea can be produced via hydrolysis of arginine into urea and ornithine or via an amphibian-like uricolytic pathway, that successively degrades uric acid into allantoin, allantoic acid and finally urea and glyoxylic acid (Figure 5)^25^. Here, we identified urea, allantoin and glyoxylate in larvae, suggesting the uricolytic pathway as the primary route for larval urea production. These metabolites were depleted in germ-free larvae, indicating a potential reduction in nitrogen excretion in these conditions, which may reflect a reduction in dietary acquisition of amino acids and nutrients in the absence of bacteria. In line with this hypothesis, the growth of larvae is slowed down after bacterial decolonisation, with a concurrent downregulation of *hexamerin* genes^4^. Hexamerins are haemolymph proteins involved in the transport and storage of amino acids in preparation for metamorphosis and thus their abundance possibly reflects the level of amino acid storage and ammonia detoxification via urea.

Free amino acids were however not particularly enriched in colonised larvae, while in blood-fed adults, amounts of amino acids and of excretion metabolites positively correlate^23^. While these differences may reflect metabolic distinctions between larvae and blood-fed adults, the microbiota impact on amino acid levels in blood-fed adults might be unrelated to nitrogen excretion. The lack of amino acids in blood-fed adults treated with antibiotics may be linked to a delay in blood digestion. In line with this interpretation, methionine which is present at the amino terminal end of proteins, was particularly enriched in mosquitoes with a conventional microbiota. In larvae and adults, pathways involved in nitrogen excretion are positively and negatively affected by the microbiota respectively, which may be due to differences in nitrogen richness in larval food vs blood. Another difference between studies is the antibiotic treatment used on adult guts while the current germ-free model avoids antibiotic use, so further comparison based on gnotobiotic adults may disentangle treatment- and developmental stage-specific impacts.

Besides the role of the microbiota to promote nitrogen harvesting during digestion, the yeast *Rhodotorula mucilaginosa* has been shown to promote nitrogen waste recycling in *Aedes albopictus* larvae^26^. This requires degradation of urea into ammonia by urease, and subsequent ammonia incorporation into glutamate to produce glutamine. The genome of *E. coli* however does not encompass urease, so such recycling effects are not predicted to happen in our model.

Our metabolomic dataset also shows modulation of several fatty acids, lipids, and sterols by the microbiota, though a consistent pattern remains elusive. When colonised with bacteria unable to degrade fatty acids, mosquito larvae exhibited significant developmental defects, underscoring the essential role of the microbiota in supplying fatty acids to the host and/or digesting them. In accordance with this, we and others had observed an accumulation of lipids in the gut via generic lipid staining and a depletion overall in the body when mosquitoes are deprived of their microbiota^4,9,27^. Besides a role of bacteria in the beta-oxidation of fatty acid chains, this gut accumulation may derive from the failed hydrolysation of dietary triacylglycerol into free fatty acids, which then cannot be loaded onto transporters for redistribution within the body. The reduction in several free fatty acids observed in our germ-free guts aligns with this model. It is also consistent with the microbiota influence on *lipophorin* gene expression in the guts of germ-free larvae^4,27^. In germ-free whole larvae, the accumulation of free fatty acids may reflect a lack of storage into larger molecules. However, to further elucidate lipid and fatty acid dynamics in germ-free larvae, specific lipidomic analyses of dissected guts and fat bodies would be needed.

Our data also reveal lower levels of cholesterol in germ-free guts, which raised our interest as cholesterol is a precursor of the moulting and metamorphosis hormone 20-hydroxyecdysone, besides its role as cell membrane component. This lack of cholesterol may be due to a decreased import rather than due to a stronger consumption, as *sterol carrier protein-2 (AAEL026044)*, which encodes the major cholesterol transporter, is downregulated in germ-free larvae while the first downstream enzyme is not^4,28,29^. *Cytochrome p450 CYP307A1 (AAEL009762)*, which encodes a downstream enzyme involved in ecdysone biosynthesis, is upregulated in germ-free whole larvae, potentially as a response to a lack of 20-hydroxyecdysone^28,29^. Interestingly, a link between ecdysone and fatty acid degradation has been observed in adult female mosquitoes, where silencing the ecdysone receptor reduces fatty acid degradation^30^. A causal link may thus exist between our observations, where bacteria would stimulate cholesterol harvest and ecdysone metabolism, which would in turn promote fatty acid degradation; further work would however be required to test this hypothesis. We only observed a minor, marginally significant increase in development to adulthood upon cholesterol supplementation to germ-free larvae. Like all holometabolous insects, mosquito larvae must reach a critical weight before initiating metamorphosis^31^. Therefore, although cholesterol supplementation may have provided sufficient metabolites for ecdysone synthesis, germ-free larvae likely lacked other essential nutrients required to attain the critical weight necessary for the commitment for metamorphosis.

In recent years, research has highlighted the crucial role of mosquito microbiota in supplying essential B vitamins, critical for developmental success. These vitamins, particularly riboflavin and folate, play pivotal roles in various metabolic pathways crucial for larval development^12,32^. Although vitamins were below level of quantification in our metabolome analysis, the absence of a microbiota significantly perturbs metabolites involved in several pathways acting downstream and upstream of these vitamins, including oxidative phosphorylation, TCA cycle, catabolism of fatty acids, and amino acid and nucleotide synthesis.

In conclusion, while previous data on the role of bacteria in mosquito development primarily focused on vitamin biosynthesis, the use of metabolomics highlighted other aspects of such support. We identified TCA cycle and urea synthesis as primary pathways affected in mosquito larvae by the absence of a microbiota and detected additional perturbations to lipid metabolism, specifically in cholesterol and fatty acid levels and distribution. Cholesterol was present in reduced amounts in germ-free guts and our supplementation experiments suggest that it strengthens development to adulthood in germ-free conditions, in line with its essential role in moulting hormone production. Finally, bacteria-mediated fatty acid degradation appears important for development of immature mosquitoes.

## Materials and Methods

### Ethics statement

No vertebrate animal was used in the experiments reported in this article. Anaesthetized mice are sometimes used for maintenance of the colony, under the protocol # 973021 validated by ethical board # 089 of the French Direction générale de la recherche et de l’innovation.

### Bacteria strains and mosquito lines

*E. coli* HS (wild-type, WT) was grown in lysogeny broth (LB). *E. coli* HA416 (auxotroph, AUX17,48) was grown in LB supplemented with 50 μg/mL kanamycin, 50 μg/mL meso-diaminopimelic acid and 200 μg/mL D-alanine. FBE640 and FBE765 strains were obtained by transducing the Δ*fadE*::*kanR* and Δ*fadH*::*kanR* mutations from the KEIO bank into MG1655 strain. Cultures were inoculated from single fresh colonies and incubated at 30 °C, shaking at 200 rpm for 16 h prior to centrifugation to set up gnotobiology experiments. They were spun down and bacterial pellets were diluted 5 times (high dose) or 500 times (low dose) in sterile milliQ water.

The *Ae. aegypti* New Orleans colony was maintained under standard insectary conditions at 28–30 °C on a 12:12 h light/dark cycle. Gnotobiotic mosquitoes were maintained in a climatic chamber at 80 % relative humidity on a 12:12 h light/dark 30 °C/25 °C cycle.

### Gnotobiology procedures and sample collection

Gnotobiotic mosquitoes were generated following previously described experimental procedures^4,33^. Briefly, eggs were surface sterilized using successive 5 min washes in 70 % ethanol, 1 % bleach and 70 % ethanol and rinsed three times in water and kept in sterile milliQ water overnight. The next day, sterile larvae were individualized in 24-well plates and provided with autoclaved food (Tetramin baby®) and a suspension of HS or HA416 bacteria to produce control and sterile larvae, respectively. Two days later, larvae were inspected to detect any third instar larvae (L3). They were excluded from the experiment as their age as L3 was undetermined. Then, L3 moulted in a time window of 5 h were washed in sterile milliQ water and transferred to 24-well plates in sterile water with fresh food. The latter point is slightly different compared to our transcriptomic study^4^ where colonized larvae were not transferred. Even though such transfer temporarily disturbs colonisation in control larvae^4^, we thought that it was particularly important for metabolomic analysis that the dietary conditions were exactly the same between control and sterile conditions. Whole larvae or dissected guts were sampled 12 h and 20 h after transfer, corresponding to an early timing after HA416-colonized larvae became sterile (95 % at 12 h^4^) and a late timing shortly before L3 would moult to the fourth instar. They were kept around -40 °C in700 µL of 80 % methanol during dissection and transferred in a -80 °C freezer for a few hours, until sampling was finished, and methanol quenching was performed on the same day.

### Methanol quenching

Samples were homogenized at 9000 rpm for 2x60s in a Precellys Evolution homogenizer (Bertin Technologies) and cellular debris were spun down at 14,000 rpm for 15 min at 4 °C. After sampling the supernatant, the pellet was mixed with 700 µL of pre-cooled 80 % methanol, homogenized and centrifuged again. Both supernatants of a sample were pooled and dried in a vacuum drier (Eppendorf) at room temperature until complete solvent evaporation. They were kept at -80 °C until lyophilization and shipping with ice packs.

### Dual phase extraction and sample preparation

Samples were resuspended in 300 μL of CHCl_3_/MeOH (2:1), vortexed for 30 s and supplemented with 300 μL of water. They were spun at 13,000 rpm for 10 min at room temperature and the top aqueous layer was then transferred into an inactivated glass vial and dried in a vacuum drier. Samples were stored at −80 °C. The lower organic layers were not used in this experiment. For GC-MS, samples were derivatized by a two-step methoximation-silylation derivatization procedure. The dried samples were first methoximated using 20 μL of 20 mg/mL methoxyamine hydrochloride in anhydrous pyridine at 37 °C for 90 min. They were then silylated with 80 μL of N-methyl-N-(trimethylsilyl)trifluoroacetamide (MSTFA) at 37 °C for 30 min.

### GC-MS data acquisition

GC-MS analysis was performed on an Agilent 7890 Gas Chromatograph (Agilent, Santa Clara, CA, USA) equipped with a 30 m DB-5 ms capillary column with a 10 m DuraGuard column coupled to an Agilent 5975 MSD system (Agilent, Santa Clara, CA, USA) which operates under electron impact ionization. Samples were injected with an Agilent 7693 AutoSampler injector (Agilent, Santa Clara, CA, USA) into deactivated spitless liners, following a temperature gradient detailed in Kind et al. 2009, using helium as the carrier gas. Quality control samples were produced by pooling different samples and analysed along with the other samples. PCA plot showed a clear clustering of QC samples at the centre of all experimental samples, indicating reproducible sample analysis (Figure S3). Metabolites were identified and quantified using a workflow described by Behrends et al^34^. While the initial sample production and data acquisition were initially performed on 5 independent replicates, the first one did not include any contamination checks while the last four ones did. Hence, we decided to include only the four replicates based on contamination-free quality check. A sample of whole larvae 12h was also excluded from the analysis because of failed derivatization.

### Cholesterol supplementation

Cholesterol solutions were prepared by dissolving 100 mg of cholesterol (C75209; Sigma) in 6 mL acetone. The solution was sterilized using a 0.2 µm filter, evaporated overnight inside a biosafety cabinet, and subsequently ground into a fine powder. This powder was resuspended in 25 mL of autoclaved food suspension to obtain a 4 mg/mL cholesterol stock solution. A control solution was prepared following the same procedure, omitting cholesterol. The stock solution was then diluted 1:3, 1:30, or 1:300, and 40 µL of each dilution was administered to larvae, resulting in final concentrations of 106, 35.6, 3.6 or 0.36 µg/mL, respectively.

### Statistics and data analysis

Data were analysed using MetaboAnalyst (version 6.0)^35^. All statistical results can be found in Table S2. Prior to normalization, geometric means of all metabolite peak intensities were calculated for each sample and conditions were compared with two-sided paired t-test. Then, a total peak intensity normalization was performed for each sample, to account for differences in amount of material input. Data were normalized by square root transformation and auto scaling. Quality Log2-transformed ratios between germ-free and colonized conditions were calculated for each type of sample at each time point and each replicate and analysed by t-test for analyses and Volcano plots. Paired ratios were calculated and a log_2_ transformation was applied. For volcano plots and KEGG pathway enrichment analysis, thresholds were set at *p* = 0.1 maximum and |log_2_(FC)|= 0.3 minimum (30% enriched). Metabolomic data are very variable; hence, no correction was applied to this t-test. This statistical analysis (as any) thus needs to be considered with caution, but this strategy allowed us to keep a sufficient number of metabolites of interest to look at pathway enrichment. As many metabolites are part of several pathways, we consider that a pathway’s enrichment can be determined if it contained at least of four metabolites detected in the dataset (Table S1). Metabolomic pathways were scrutinized using the Kyoto Encyclopedia of Genes and Genomes (KEGG). Analysis of developmental success and CFU counts were performed with generalised linear mixed models (GLMM) or a linear regression (lmer) using the lme4, lmerTest and lsmeans packages in R (version 4.3.0). For developmental success data, an ANOVA was performed on a logistic regression (glmer), while for CFU counts an ANOVA was performed on a linear regression (lmer). The replicate was set as a random effect in both cases.

## Acknowledgements

We thank Siegfried Hapfelmeier for provision of HS and HA416 bacteria, Johan Claes Schönbeck for technical help and Yannick Estevez for sample lyophilization. We are grateful to late Jean Issaly for mosquito maintenance.

## Funding

This work has been supported by LabEx IBEID (The Laboratory of Excellence (LabEx) Integrative Biology of Emerging Infectious Diseases—grant no. ANR-10-LABX-62-IBEID) and ANR JCJC MosMi to MG (grant no. ANR-18-CE15-0007).

**Figure S1. Metabolites significantly affected by bacterial colonization in gut and larval samples at different time points.** Volcano plots showing differential metabolite abundances between colonized and germ-free conditions in gut (**A**-**B**) and whole-larva (**C**-**D**) samples collected at 12 h (**A**,**C**) or 20 h (**B**,**D**) post bacterial decolonization. Metabolites enriched in colonized samples are indicated by orange arrows; those enriched in germ-free samples by black arrows. A fold change threshold of 1.2 and a t-test *p* value threshold of 0.1 were applied. Both fold changes and *p* values are log-transformed for visualization.

**Figure S2. *fadE* and *fadH* mutants are not deficient in colonizing breeding water and larvae and not pathogenic to larvae.** (**A**), CFUs of wild type (WT, khaki), Δf*adE* (purple) or Δ*fadH* (green) *E. coli* in larval breeding water at the time of colonization, 1 day and 2 days later. (**B**) CFU quantification of WT (white), Δf*adE* (purple) or Δ*fadH* (green) *E. coli* in fourth-instar larvae. Statistical significance on the CFUs was assessed using a linear model with bacterial strain and CFUs as fixed factors and the replicate as a random factor. (**C**) Development of larvae when providing a high quantity of bacteria at the start of the experiment (10^8^ CFUs/mL). Bar plots show the proportions of adults (black, purple or green) and developmentally blocked or dead larvae (grey). Statistical significance on the proportion of adults was assessed using a generalized linear mixed model (GLMM) using the proportion of adults and the cholesterol concentration as fixed factors, and the replicate as a random factor. All plots show results from five independent replicates. The number of individuals analysed in each replicate is indicated below the plots.

**Figure S3. Principal Component Analysis (PCA) of gut and larval metabolomes and quality control (QC) samples.** PCA plot showing the distribution of experimental and QC sample along the first five principal components. Sample type is colour-coded: colonized (orange), germ-free (white), and QC (green). Time points are indicated by symbol shape: circles for 12 h, squares for 20 h, and crosses for QC samples.

## Supplementary table legends

**Table S1. List of metabolites and their raw peak intensities.** For each metabolite, its general category, retention time (RT) and intensity in each sample is indicated. Values in grey belongs to a replicate that was excluded because lacking appropriate contamination checks. Values in red belong to the sample that did not derivatize. QC: quality control.

**Table S2. Statistical analysis results.** For Figure 2, PERMANOVA F, R-squared and *p* value results are indicated for each discriminatory variable (tissue, time, colonization, time, colonization*time). Significant *p* values are highlighted in red.

For Figures 4 and S1, the fold change values (and their the log_2_ transformations) and the raw t-test *p* values (and their -log_10_ transformations) are shown for the metabolites with *p* < 0.1 and -1.2 < fold change < 1.2. Metabolites significantly affected by bacterial decolonization in both 12 h and 20 h time-points are highlighted in green.

For Figure 7, GLMM results are shown for models using the proportion of blocked larvae, dead larvae, dead pupae and adults, and the cholesterol concentration as fixed factors, and the replicate as a random factor.

For Figure S2, LMM and GLMM results are showed for models using the number of CFUs in water (Figure S2A) or larvae (Figure S2B) or the proportion of adults (figure S2C) and the bacterial strain as fixed factors, and the replicate as a random factor.

